# Habitat loss causes long transients in small trophic chains

**DOI:** 10.1101/2020.05.15.098863

**Authors:** Blai Vidiella, Ernest Fontich, Sergi Valverde, Josep Sardanyés

## Abstract

Transients in ecology are extremely important since they determine how equilibria are approached. The debate on the dynamic stability of ecosystems has been largely focused on equilibrium states. However, since ecosystems are constantly changing due to climate conditions or to perturbations such as the climate crisis or anthropogenic actions (habitat destruction, deforestation, or defaunation), it is important to study how dynamics can proceed till equilibria. In this contribution we investigate dynamics and transient phenomena in small food chains using mathematical models. We are interested in the impact of habitat loss in ecosystems with vegetation undergoing facilitation. We provide a thorough dynamical study of a small food chain system given by three trophic levels: vegetation, herbivores, and predators. The dynamics of the vegetation alone suffers a saddle-node bifurcation, causing extremely long transients. The addition of a herbivore introduces a remarkable number of new phenomena. Specifically, we show that, apart from the saddle node involving the extinction of the full system, a transcritical and a supercritical Hopf-Andronov bifurcation allow for the coexistence of vegetation and herbivores via non-oscillatory and oscillatory dynamics, respectively. Furthermore, a global transition given by a heteroclinic bifurcation is also shown to cause a full extinction. The addition of a predator species to the previous systems introduces further complexity and dynamics, also allowing for the coupling of different transient phenomena such as ghost transients and transient oscillations after the heteroclinic bifurcation. Our study shows how the increase of ecological complexity via addition of new trophic levels and their associated nonlinear interactions may modify dynamics, bifurcations, and transient phenomena.

## I. INTRODUCTION

Ecosystems are highly nonlinear, complex dynamical systems [1–5]. Nonlinearities in ecology, driven by density-dependent interactions such as competition, cooperation, or victim-exploiter dynamics, introduce far from trivial cause-effects in the population dynamics. This is of special importance when dealing with ecosystems’ responses to perturbations, both natural or of anthropogenic origin i.e. deforestation, habitat destruction, animal hunting. Nonlinear dynamics in ecosystems include intrinsic oscillations [6, 7] or deterministic chaos. For instance, chaos has been suggested to be found in vertebrate populations [8–12], plankton dynamics [13, 14], and insect species [15–18].

Nonlinearities also give place to TIPPING POINTS, which have important consequences in species persistence and extinctions. In the last years, tipping points are becoming extremely relevant in ecology, due to the anthropogenic effects on the ecosystem (i.e. climate change). Not only individual ecosystems can develop an abrupt shift (from glass melting to cyanoblooms) [19, 20], the whole planetary biome may change [21–25]. One example of this abrupt change is the transition between the past vegetated Sahara to the current desert [26, 27]. In these ecosystems, the aridity restricts the availability to water, narrowing the vegetation cover. Moreover, external pressures like grazing or deforestation have shown that for the same aridity levels two possibilities are possible, vegetated or desert [28, 29]. In lower levels of aridity multiple ecosystems can be observed (savanna, forest, grasslands, and shrublands) but for some levels of aridity abrupt changes occur [30]. Furthermore, in the framework of systems biology, a tipping point driving populations to extinction has been recently described experimentally in yeast [31].

Tipping points can be studied by means of bifurcation theory (Box 1). It is important to note that long transients typically arise near bifurcations, where slowing down phenomena take place. Of particular interest are the so-called SADDLE REMNANT or GHOSTS, suggested as transient-generator mechanisms in ecological systems in Ref. [32]. Ghosts arise in biological dynamical systems with strong feedbacks such as cooperation [33–35] and facilitation processes [36, 37] (see below). Other suggested mechanisms responsible for transients are chaotic saddles responsible for transient chaos [38, 39], spatial systems [40], linear systems with varying time scales, coupled oscillators, and stochasticity [41] (see also Refs. [42–44]).

### Box 1: Bifurcations and their transients

Bifurcations involve qualitative changes in dynamics when one or more parameters are changed [60, 112]. They can be generically classified as local or global. Local bifurcations involve fixed points (stability changes, collisions). Global bifurcations involve transitions associated to bigger objects such as global manifolds, limit cycles or STRANGE ATTRACTORS. Bifurcations are key phenomena in transients since they typically involve long delays of the orbits when parameters are close to bifurcation values. The bifurcations identified in this article are explained as follows.

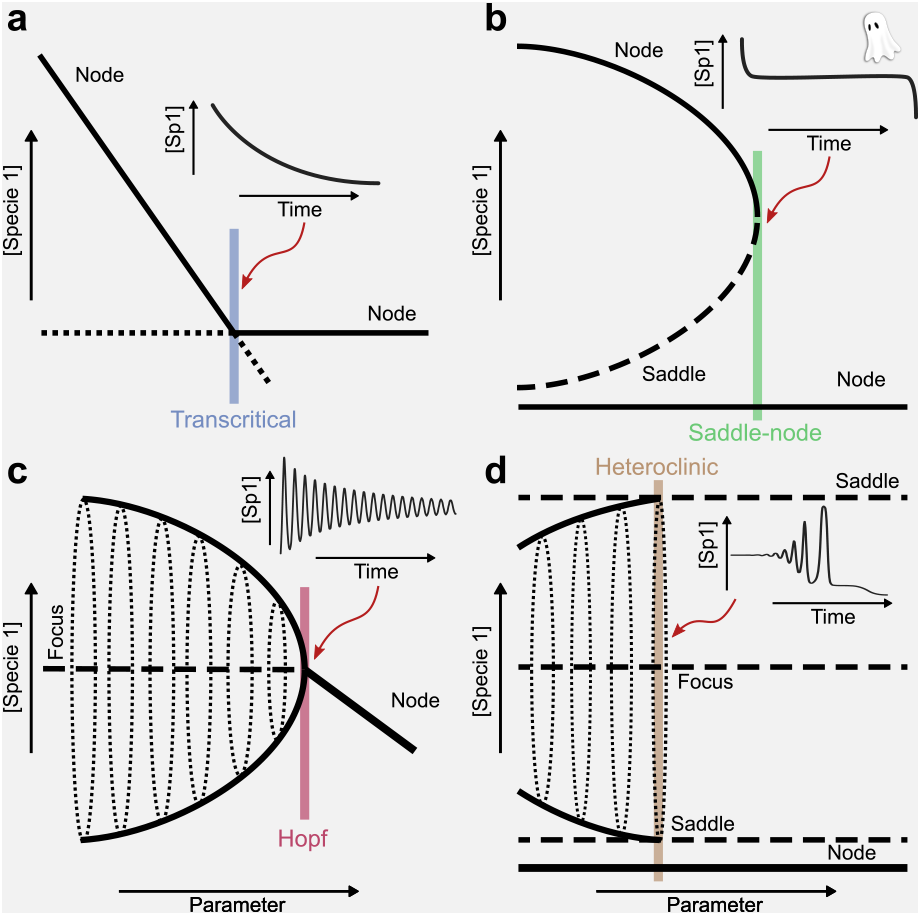

**Local bifurcations**:

a. *Transcritical:* collision (without destruction) of two fixed points with an interchange of stability between them.
b. *Saddle-Node:* basic mechanism of creation or destruction of equilibrium points (saddle point-node). After the collision a saddle remnant (ghost) continues influencing the dynamics, causing extremely long delays, which follow the inverse square-root scaling law [32–36]. Bifurcation identified in yeast experiments [31], in an electronic circuit (for this case the scaling law was also identified) [61], and in elastic structures [68].
c. *Supercritical Andronov Hopf:* transition from a stable FOCUS to an unstable one spiralling outwards towards a stable cycle. Bifurcation identified in convection [69] and chemical reactions [70, 71]. Slow passage close to bifurcation was also identified in Ref. [71]. **Global bifurcations:**
d. *Heteroclinic:* the global manifolds of two saddle points collide (at bifurcation both equilibria have a heteroclinic connection). Transients with large-amplitude oscillations.

Recent research shows how human activity could be pushing ecosystems towards tipping points [45, 46]. Habitat fragmentation is one of the major causes of species extinctions [47–50]. The loss of wild areas [51] is increasing in the last years, also making species extinctions more and more frequent, leading to the so-called sixth mass extinction [52–56]. This, among other anthropogenic alterations, can lead to run-away effects from the bio-geo-physical planetary processes [19, 20, 25] e.g. when grass disappears the soil is exposed to wind erosion, impairing the establishment of grass again [30]. In this sense, habitat destruction plays a major role in determining the fate of ecosystems. Figure 1(a) shows the planetary distribution of fragmented habitats, which comprise the most developed countries as well as the currently developing ones. Several works have studied the impact of habitat destruction with simple models [37, 50, 57], giving clues on the expected dynamics and bifurcations.

**FIG. 1:**
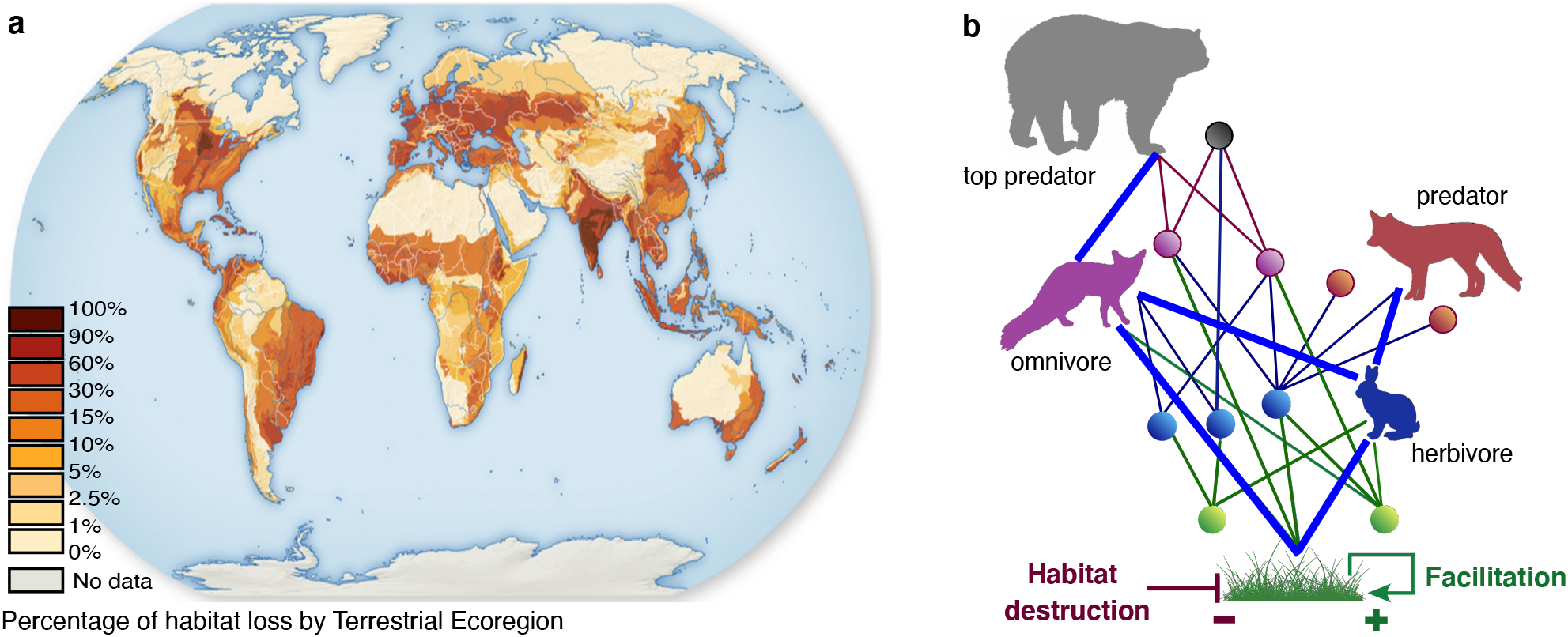
(a) Habitat loss is an important ecological handicap for species persistence. Here we display the percentages of habitat loss divided in terrestrial eco-regions (source: http://habitatlossfragmentation.weebly.com/habitat-loss.html). (b) Trophic networks underly ecosystem dynamics. Here we display a large trophic network with different trophic levels: primary producers (green); herbivores (blue); omnivores (violet); predators (brown and grey). In this manuscript we will address the subnetwork highlighted by blue thick links by means of a simple dynamical model, focusing on the interactions vegetation-herbivore-predator and considering facilitation processes among primary produces and habitat destruction. Finally, the model model adding an omnivore and a top predator will be introduced. Our main goal is to characterise how an increase in ecological complexity (addition of different trophic levels) impacts on the bifurcations and the transients.

### Glossary

**Tipping point:** qualitative change in dynamics produced by changes in parameter/s (equivalent to bifurcation).

**Equilibrium or fixed point:** state of the dynamical system that does not move as time changes. It satisfies *dx/dt* = 0 for a single variable system. It can be either stable or unstable.

**Phase space:** set of all possible states of the dynamical system. This space is built using the dynamical variables as axes.

**Eigenvalues of a fixed point:** factors of contraction or expansion along their associated invariant directions (generated by the corresponding eigenvectors) of the linearized system at the fixed point. They are used to determine the stability of equilibrium points.

**Orbit:** path described by the dynamical variables in the phase space as time changes.

**Saddle-node remnant (ghost):** effect found close to a saddle-node bifurcation after the equilibrium points have disapeared which causes extremely long transients [32, 33, 35, 60]. Identified in electronic circuits [61].

**Delayed transition:** slowing down for saddle-node bifurcations casued by the ghost.

**Monostability:** dynamical system with a single stable equilibrium.

**Bistability:** dynamical system with two stable equilibria. Classic ecological examples are shallow and eutrophic lakes [62, 63], vegetated or desert states in semi-arid ecosystems[28, 30] (see also [64]).

**Globally asymptotically stable equilibrium point:** equilibrium point that is stable and such that all orbits of the system converge to it.

**Focus:** equilibirum point for which all nearby orbits describe a spiral behaviour, either stable or unstable.

**Stable limit cycle:**periodic oscillation which attracts all nearby orbits.

**Strange attractors:** attracting objects in phase space with complicated geometric structure (fractal). Usually the dynamics on them is chaotic.

**Stable manifold of an equilibrium:** set of points that converge to the equilibrium, usually having less dimension than the one of the system. In the plane they are curves that separate regions with different dynamical behavior.

**Unstable manifold:** same as the stable manifold but with backward time.

**Heteroclinic connection:** set of orbits connecting two different equilibrium points of saddle type.

In this article we investigate a simple mathematical model describing the dynamics of a small trophic chain including facilitation at the level of primary producers and habitat destruction/loss. Our interest is to see how increasing ecological complexity i.e., the addition of trophic levels and interactions, affects dynamics, focusing on bifurcations and transients. The interplay between the triad *habitat loss-facilitation-trophic relations* has not been investigated in detail, especially at the level of transients. Facilitation (positive pairwise interactions between individuals leading to the benefit of at least one of the interacting partners[58]) is a key ecological process in many ecosystems e.g. drylands. This positive effect can occur either directly or indirectly. An example of the former includes shading mechanisms that reduce water or nutrient stress. The later would include removing competitors or deterring predators [59].

To ease the reading to an audience non familiarised with technical mathematical details, we provide a glossary of words and a box gathering the main characteristics of the bifurcations identified in our work.

## II. MATHEMATICAL MODEL AND RESULTS

### Model 1. Vegetation with facilitation

Starting with the simplest system we aim at characterising how the increase in ecological interactions and the addition of species impact dynamics, bifurcations, and, especially, transients. The first model describes a population of primary producers (e.g., plants, cyanobacteria, algae) with facilitated reproduction and intra-specific competition, together with habitat destruction and density-independent death rate. Facilitation is introduced with the nonlinear term *αV*^2^, *α* being the intrinsic growth rate of plants. This growth kinetics has been used to model autocatalysis [32] and ecological facilitation [37], in which a replicator enhances its own growth. As a difference from exponential growth, *αV*, the autocatalytic one is known to produce hyperbolic dynamics, in which an infinite population can be achieved in finite time. This can be seen from the time solution for 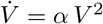, given by *V*(*t*) = *V*(0)/(1 − *V*(0)*αt*).

In our approach, the reproduction of the vegetation is constrained by a logistic-like function 1 − (*D* + *V*)/*C*_0_, where constant *D* is the fraction of habitat destroyed (see Refs. [37, 50, 57, 65, 66]) and *C*_0_ is the carrying capacity (set to 1 for simplicity). Hence, this first model considers four main ecological processes: (i) reproduction of plants with facilitation proportional to parameter *α* > 0, (ii) habitat destruction modelled with parameter *D* ∈ [0,1], (iii) intraspecific competition, and (iv) density-independent death rate (*ϵ*) of plants. Parameter *D* takes values between 0 and 1 since it represents the (normalised) fraction of habitat destroyed. The model is given by:

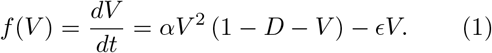

The previous model has been recently studied in Ref. [37], and will be briefly discussed below. Let us first study the simplest model for vegetation setting *ϵ* = 0. This system has two EQUILIBRIUM or FIXED POINTS, obtained from *ẋ* = 0, with 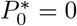 and 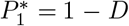. The local stability of these equilibria can be computed from λ = *df*(*V*)/*dV* = 2*αV*(1 − *D* − 3/2 *V*). It is easy to see that 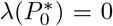 and that 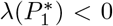 under the assumption that 0 < *D* < 1, thus 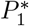 being stable within this range of *D*. This can be seen in Fig. 2(a), where the dynamics of Eq.(1) have been computed numerically[114] setting *ϵ* = 0. The population declines linearly as *D* increases. The stability of this equilibrium, computed from 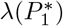 is indicated with a red line, which is equal to zero at *D* = 1.

**FIG. 2:**
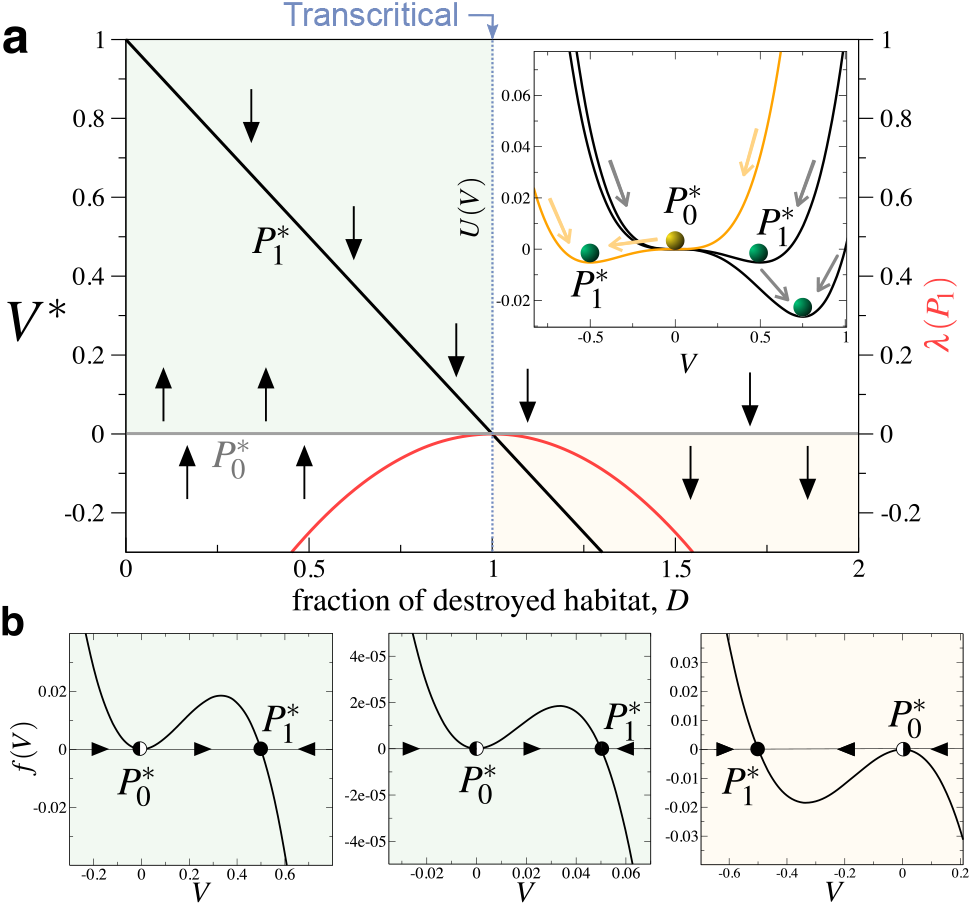
Dynamics of Eq.(1) with *ϵ* = 0. (a) Equilibrium values of the vegetation at increasing the fraction of habitat destroyed, *D*, given by 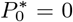 and 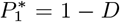. The green rectangle displays the biologically-meaningful values of *D*, with 0 ≤ *D* ≤ 1. Notice that within this range the fixed point 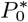 is unstable (small vertical arrows denote the stability). Also, 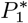 is always stable except at the bifurcation value *D* = 1, where 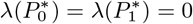 (the red curve shows the value of 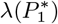). This means that transients towards e.g. 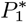, become extremely long as *D* → 1. The inset in (a) displays the potential function (2) computed with *D* = 0.25 and *D* = 0.5 (black lines), and *D* = 1.5 (orange line). Notice that 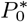 is half-stable: repeller for *V*(0) > 0 and 0 ≤ *D* < 1. At *D* = 1 this stability for 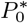 is changed. (b) Flows on the line displayed in the space (*f*(*V*), *V*): (left) *D* = 0.25; (middle) *D* = 0.95; (right) *D* = 1.5 (here black dots indicate stable equilibria while black-white dots denote half-stable points). In all panels we have set *α* = 1.

The case 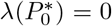 does not provide any information on the stability of the origin. To determine the stability of this equilibrium we will compute a potential function and use graphical methods. A useful way to study and visualise the stability of the fixed points in a one-variable system is by means of the potential function, *U*(*V*), given by:

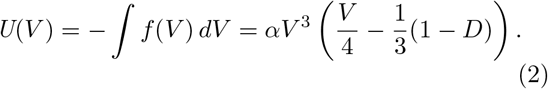

The inset in Fig. 2(a) displays the potential, which has a minimum at 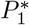. Note that the origin is semistable. This can be clearly seen plotting *f*(*V*) against *V* (Fig. 2(b)). Due to the simplicity of this model one can compute the transient times to the fixed point 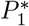 analytically, from *dV*/*dt* = *f*(*V*), using:

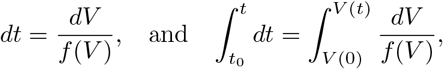

obtaining, taking *t*_0_ = 0:

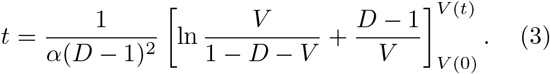

From the previous expression it is easy to see that *t* → ∞ as *D* tends to the bifurcation value.

As we previously mentioned, the dynamics of Eq. (1) with *ϵ* > 0 has been recently investigated [37]. This system has two different dynamical regimes (MONOSTABILITY or BISTABILITY) as a function of the model parameters. This model has three equilibrium point, namely 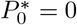 and the pair

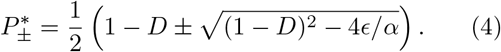

Notice that the pair 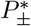 will be biologically meaningful whenever the discriminant of the fixed points (4) is either positive or zero. At the bifurcation value the discriminant is zero and these two fixed points collide in a saddle-node bifurcation. This occurs at

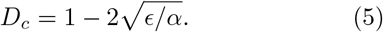

Linear stability analysis for this system indicates that when *D* < *D_c_* both 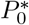 and 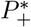 are locally asymptotically stable, while 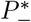 is a repeller. At *D* = *D_c_*, 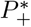 and 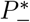 collide and are destroyed, and thus the only remaining equilibrium is 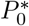, which becomes GLOBALLY ASYMPTOTICALLY STABLE [37]. The bifurcation diagram using *D* as control parameter is shown in Fig. 3(a). It is known that just after a saddle-node bifurcation, transients experience an extremely long delay, called a DELAYED TRANSITION [60, 67]. This is due to a SADDLE REMNANT or GHOST that attracts the orbits towards the region of the PHASE SPACE where the collision occurred, although no fixed points are present. This is why this is called a ghost, phenomenon that was claimed as a transient generator mechanism in ecological systems in Ref. [32]. It is known that the passage time, *t_p_*, through the ghost follows a universal scaling law of the form *t_p_* ~ (*D* − *D_c_*)^-1/2^, as panel (b) in Fig. 3 shows. This ghost can be easily observed in time series, where trajectories settle onto a flat, extremely long bottleneck before a rapid collapse (Fig. 3(d)).

**FIG. 3:**
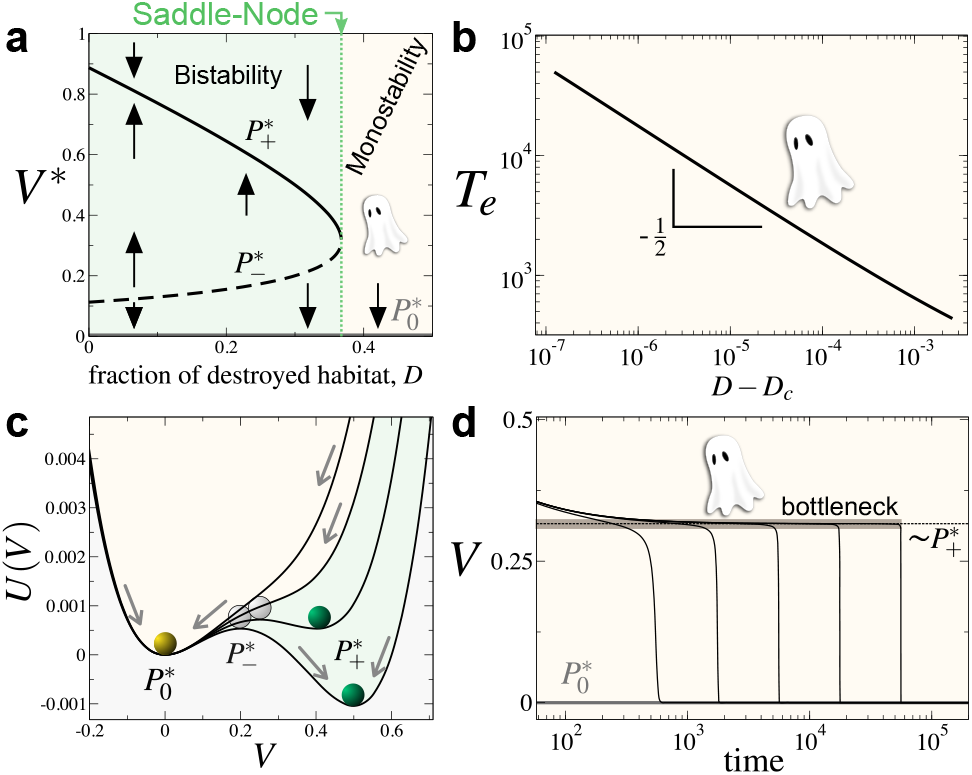
Dynamics of Eq. (1) with *ϵ* > 0. (a) Saddle-node bifurcation, which leaves a ghost after bifurcation threshold and the system becomes monostable (yellow region). (b) Inverse square-root scaling law tied to the extinction transients trapped by the ghost. (c) Potentials for *D* = 0.3, *D* = 0.35, *D* = 0.4, and *D* = 0.45. In all panels we have used *α* = 1 and *ϵ* = 0.1. (d) Delayed transitions on the ghost bottleneck with *D* = *D_c_* + *ω*, with (from left to right): *ω* = 10^−3^, *ω* = 10^−4^, *ω* = 10^−5^, *ω* = 10^−6^, *ω* = 10^−7^.

A potential function can be also computed for this system, now having:

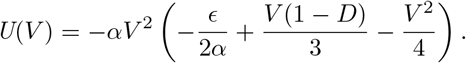

The potential is displayed in Fig. 3(c) for several values of *D*. For *D* < *D_c_* (solid lines) two wells are found, corresponding to the two stable states (*P*0* and 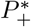) resulting in bistability, that will be achieved depending on the initial conditions. Evidences of multiple states have been recently identified in semi-arid ecosystems [28, 30]. Once the fraction of habitat destroyed surpasses its critical value *D_c_*, a single well is found. This single stable state is given by the equilibrium point 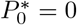, which involves extinction.

### Model 2. Vegetation-herbivore system

In this section we consider another trophic level by adding a herbivore species to the dynamics of vegetation. That is, considering model (1) plus the new species, *S*_1_. The model now reads:

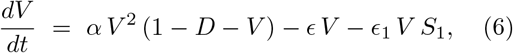

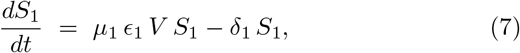

Here we consider that vegetation is consumed by the herbivore at a rate *ϵ*_1_ > 0. Also, we consider that *S*_1_ reproduces proportionally to *μ*_1_ *ϵ*_1_, where *μ*_1_ < 1 is the effective reproduction rate. For simplicity *μ*_1_ and *δ*_1_ will be labeled as *μ* and *δ*. The equilibrium points of Eqs. (6)–(7) are:

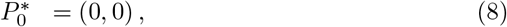

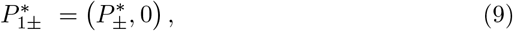

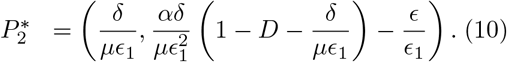

Note that equilibrium 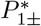 involves the extinction of the herbivore species and the persistence of vegetation, with same equilibria as the ones given in Eq. (4). Equilibrium 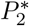 involves coexistence. This equilibrium will be biologically-meaningful i.e. positive, when the fraction of habitat destruction is below the critical value:

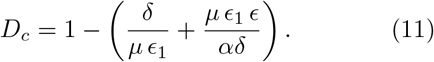

The stability of the equilibrum points is computed from the linearised system given by the Jacobian matrix:

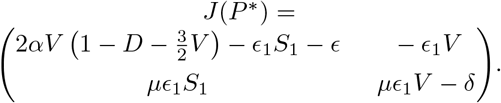

The stability of an equilibrium 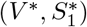 is computed from the sign of the EIGENVALUES obtained from the characteristic equation 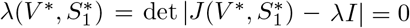, *I* being the identity matrix. The eigenvalues of 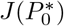 are λ_1_ = −*ϵ*, λ_2_ = −*δ*, meaning that the origin is always a local attractor. The Jacobian matrix evaluated at equilibrium 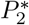 is:

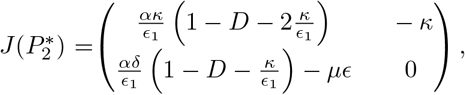

with trace

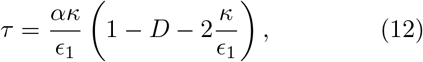

and determinant

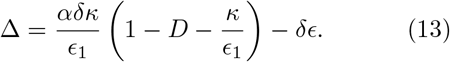

We note that Δ > 0 in the range of parameters where 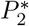 exists. The stability of this fixed point can be determined with the sign of the trace, being an attractor when *τ* < 0, and a repeller when *τ* > 0. The bifurcation value 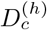, involving the SUPERCRTICAL Andronov Hopf bifurcation, is here given by:

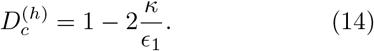

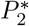 is stable when 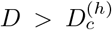. Since Δ > 0 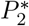, is a focus. According to the previous calculations, the following local bifurcations are found in Eqs. (6)–(7) (see Fig. 4 and Box 1):

1. Saddle-node bifurcation identified in Section 2.1., causing delayed transitions (ghost transients).
2. Supercritical Andronov Hopf bifurcation when 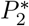 changes its stability from a stable focus to an unstable one (condition given by Eq. (14)). This will also induce long transients towards the oscillating coexistence.
3. Transcritical bifurcation when 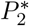 and 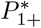 collide, occurring when *D* = *D_c_* (see Eq. (11)). This will cause a critical slowing down.

**FIG. 4:**
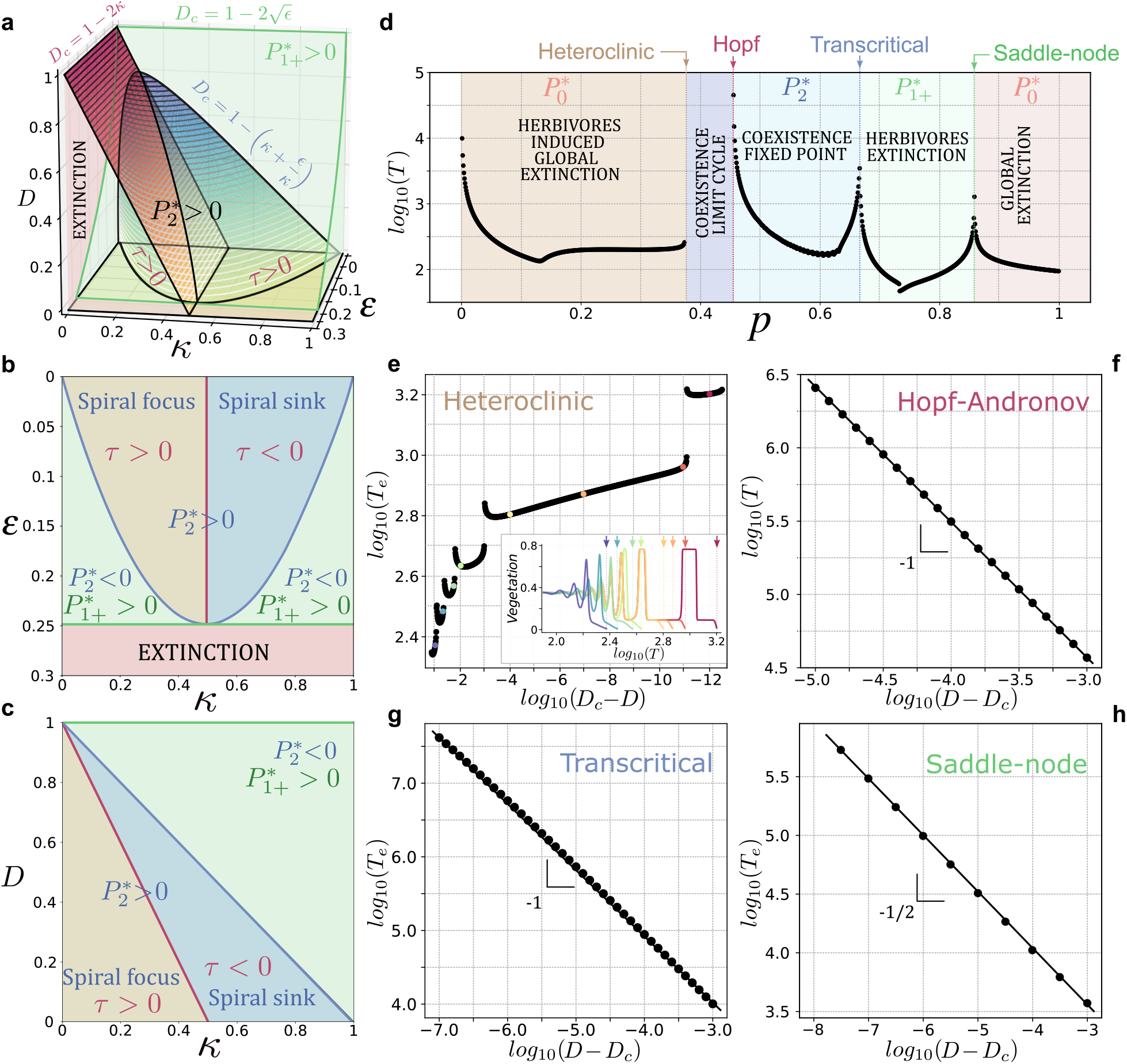
Stability conditions, bifurcations, and transients of Eqs. (6)–(7). (a) Parameter space built with (*ϵ, κ, D*) showing the existence regions of equilibria and the boundaries involving bifurcations: under the green surface *P*_1±_ exist, under the blue one 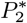 exists and their eigenvalues have imaginary part (Δ > 0). The red surface indicates the change in 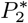 stability (being a repeller below the surface). (b) Stability regions found in the plane *D* = 0 (without habitat destructed). (c) Transitions occurring in the plane *ϵ* = 0 (without density-independent death of vegetation). (d) Times needed to achieve an attractor (given by equilibria within each coloured region) tuning simultaneously parameters (*ϵ, κ, D*) by using *p* × (0.2*ϵ, κ*, 0.2*D*), with *p* ∈ [0,1]. Panels (e) to (h) display how transients change as the bifurcation parameter *D* approaches to the bifurcation value *D_c_*. (e) Transients close to the heteroclinic bifurcation. The inset displays several time series for different values of *D* (indicated with the coloured dots).(f) Transient times before the SUPERCRITICAL HOF-ANDRONOV bifurcation of 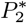 (red surface in the panel (a) and red lines in panels (b)-(c). (g) Transients arising after the the transcritical bifurcation of 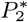 and 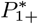 (blue surface in the panels (a)-(c). (h) Transients just after the saddle-node bifurcation giving place to the inverse square-root scaling law (green surface in panel (a)). In all the analyses *α* = *ϵ*_1_ = 1.

Equations (6)–(7) also suffer a global bifurcation given by a heteroclinic bifurcation originated from a heteroclinic connection between equilibria 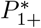 and 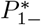. This bifurcation occurs via the collision of the global manifolds of both fixed points (see Section IV A in the *Appendix* for detailed information). Close to the bifurcation, the geometric structure of the manifold forces all the trajectories to pass very close to the fixed points 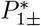 before achieving the origin, approaching the ORBITS to regions where the vector field is ex-tremely small thus causing delays and long transients. This actually causes a kind of slow-fast transient dynamics.

The dependence of dynamics on parameters and the bifurcations found in Eqs. (6)–(7) are summarised in Fig. 4: panels (a) shows the boundaries between the different dynamical regimes in the parameter space (*ϵ, κ, D*). Panels (b) and (c) display cuts in the threedimensional parameter space showing the stability of the equilibrium points and their existence. All of the bifurcations identified in the model are displayed in Fig. 4(d) tuning an auxiliary parameter *p* ∈ [0,1], allowing to cross the parameter space in (a) with a straight line by means of *p* · (0.2*e*, *κ*, 0.2*δ*), which is plotted against the times needed to achieve an attractor. Notice that transients increase near bifurcation values since the vector field tends to zero. It is known that the length of transients close to local bifurcations typically follow scaling relations between the parametric distance to the bifurcation value [60]. As we previously discussed, this scaling for saddle-node bifurcations is given by the inverse square-root scaling law [60].

These delays, known as delayed transitions for saddle-node bifurcations, are generically named as critical slowing down. Different exponents for these scaling laws are known to exist, being − 1/2 for the saddle-node bifurcation and −1 for the transcritical bifurcation (see Fig. 4(h,g)). The supercritical Andronov Hopf bifurcation also displays a scaling exponent of −1 before the emergence of the STABLE LIMIT CYCLE, as shown in Fig. 4(f). As mentioned, the heteroclinic bifurcation is a global bifurcation, and involves changes in the global invariant manifolds, while fixed points neither change (remarkably) their stability nor collide. Figure 4(e) shows that, as the critical value of the heteroclinic bifurcation 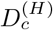 is approached, the time to extinction *T_e_* increases in a discontinuous way (the computation of 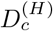 can be found in Section IVB in the Appendix). These jumps are tied to the emergence of new cycles as *D*^(*H*)^ → *D_c_*. This can be clearly seen in Fig. 5. Specifically, Fig. 5(a) shows a time series extremely close to the bifurcation value, where the dynamics undergoes 11 cycles[115] For some lower *D* values the number of cycles diminishes, providing discontinuities in times as shown in Fig. 5(b) (see also Fig. 4(e)). Figure 5(c) displays how the cycles appear and accumulate at increasing *D*.

**FIG. 5:**
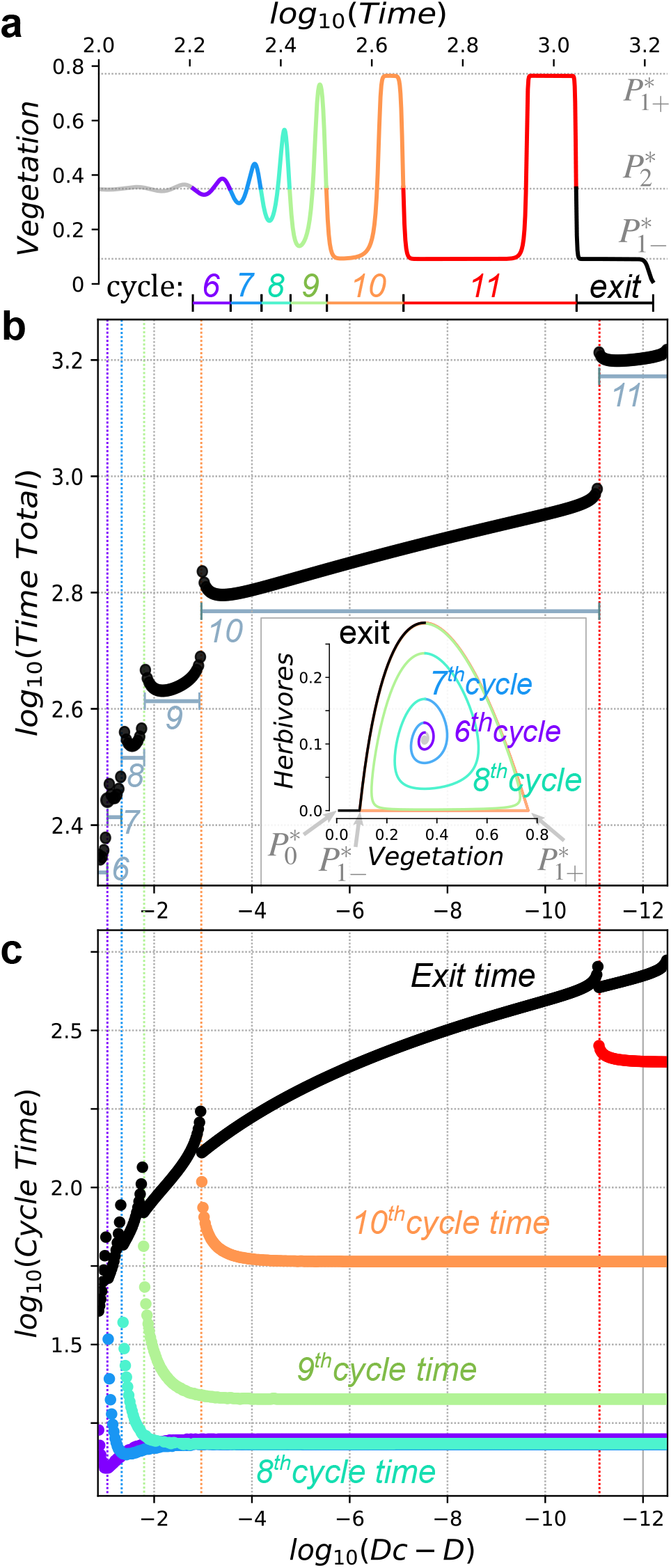
Transients close to the heteroclinic bifurcation. (a) Extinction transients after the heteroclinic bifurcation with *D_c_* − *D* ~ 10^−12^. Each colour indicates a cycle around the repeller. (b) Same as in Fig. 4(e) now showing the trajectory represented in panel (a) in a phase portrait. Here the orbit gets very close to the saddle points *P*_1+_ and *P*_1−_. (c) Time per cycle depending on the distance to the bifurcation value. For these analysis we use: *α* = *μ* = *ϵ*_1_ = 1, *κ* = 0.35, *ϵ* = 0.07 and *D* = 0.1439373616781516 ≲ *D_c_*. The initial conditions are taken near to 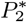.

We note that each new cycle forces the orbits to pass closer to the points 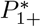 and 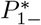, which are saddle points, and thus have stable and unstable local manifolds. This effect makes the cycles to lapse more time, giving to the time trajectories the typical square shape found in slow-fast systems (here in a transitory way, see also the inset time series in Fig. 4(e)) as well as in asymptotic heteroclinic cycles dynamics (see e.g. [3, 72, 73]). Finally, we note that the exit time (the time of residence at the left side of 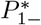) increases with the proximity to de critical parameter value. Figure 6 provides dynamics and bifurcations for the vegetation and herbivores, where each one of the local bifurcations as well as the global bifurcation can be visualised. Each new dynamical regime upon bifurcation has an associated phase portrait and time series, represented in panels (a)-(f) in Fig. 6.

**FIG. 6:**
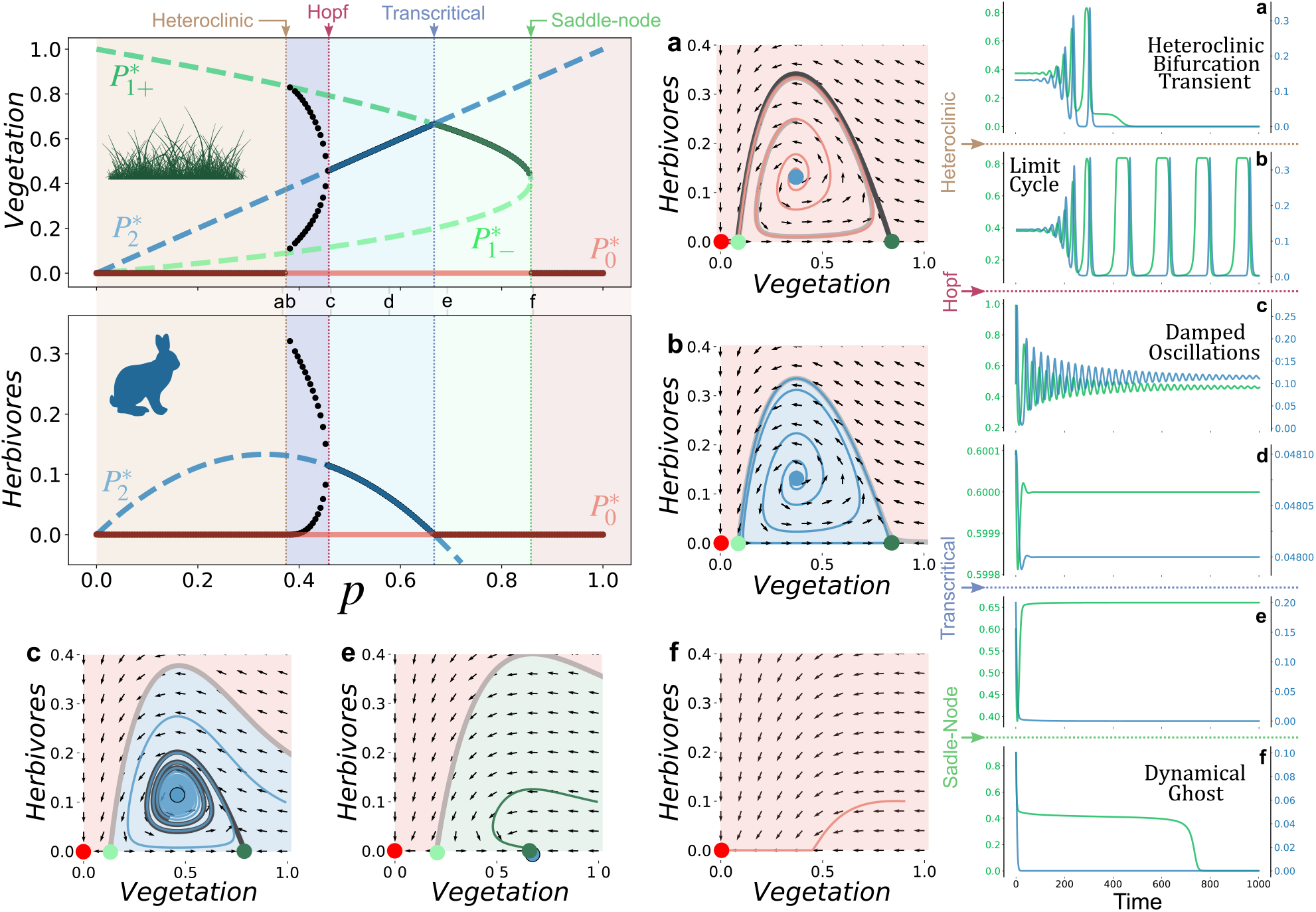
Bifurcation diagrams tuning *p* to sample the parameter space (*ϵ, κ, D*) with a straight line (more precisely, *D* = *ϵ* = 0.2*p* and *κ* = *p*). The dynamics for different regions of the bifurcation diagrams are illustrated with the small letters showing phase portraits and time series. In the first region, 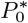 is the only stable fixed point. Just after the heteroclinic bifurcation, a long transient with a slow-fast dynamics appear (see panels (a) with *p* = 0.3735). Between the heteroclinic and the Hopf bifurcation, a limit cycle appears (panels (b. Right after the Hopf bifurcation, damped oscillations are obtained (see panels **c** with *p* = 0.46), but they get largely tenuated when the parameter distance o the transition increases (panels **d** setting *p* = 0.6). After the transcritical bifurcation the herbivores can not surviver, and vegetation remains in the ecosystem (panels (e), *p* = 0.69). Finally, when death rates are high enough, a saddle-node bifurcation occurs and all populations go to extinction (e.g. panels (f), *p* = 0.8582). Equilibrium points in the phase portraits are shown with coloured dots: 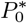 (red); 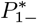 (light green); and 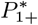 (dark green). Their basins of attraction are depicted by the coloured areas using same colours of equilibrium points. Global invariant manifolds for 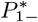 and 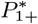 are displayed with grey and black lines in the phase portraits. In all analyses we have used *α* = *μ* = *ϵ*_1_ = 1.

Notice that all the described bifurcations, and their associated transients, can be achieved by increasing habitat destruction (*D*), increasing the death rate of primary producers (*ε*) i.e. deforestation/toxic compounds; increase of herbivores’ hunting (*δ*) increasing *κ*; or raising their effective growth rate (*μ*) lowering *κ*. One example of this last possibility, may be the reduction in the abundance of higher trophic organisms (i.e. predators). The impacts of predators is studied in the following section.

### Model 3. Vegetation-herbivore-predator system

In this section we investigate the impact of introducing a predator species, *S*_2_, on the two-dimensional, vegetation-herbivore system previously studied. The model now reads:

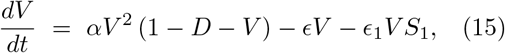

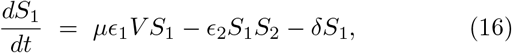

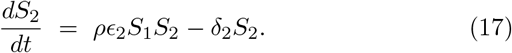

Here predators increase in numbers due to the consumption of the herbivores, *S*_1_. The predation rate is *ϵ*_2_ > 0 and the effective reproduction of predators is given by the term *ρϵ*_2_, with *ρ* < 1. Predators die at rate *δ*_2_ > 0. This dynamical system has seven equilibrium points:

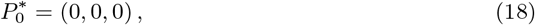

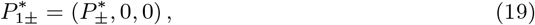

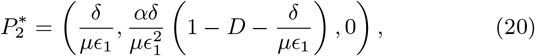

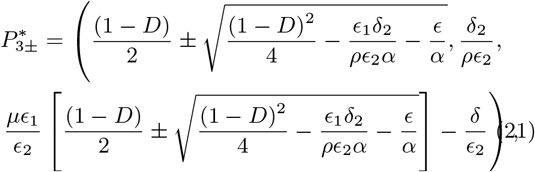

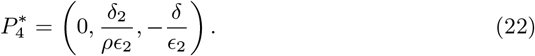

Note that, compared to the previous model, two new internal fixed points, 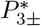, appear (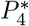 is not biologically meaningful since it has a negative component).

Equilibrium 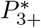 is now responsible for the coexistence of the full food chain. We note that fixed points 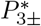 will exist i.e., 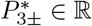, when *D* < *D_c_*, with

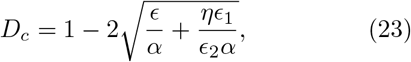

here with *η* = *δ*_2_/*ρ*. This means that equilibria 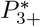 and 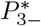 collide in a saddle-node bifurcation at *D* = *D_c_*, with 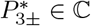 for *D* > *D_c_*.

The Jacobian matrix for this system is:

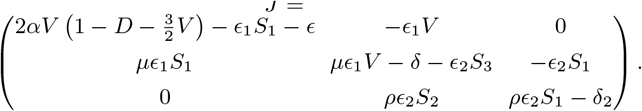

The equilibrium point placed at the origin 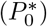 still remains as a local attractor, with eigenvalues −*ϵ*, −*δ*, −*δ*_2_. Equilibria 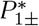, involving only the persistence of vegetation, gain a stable direction in the full phase space i.e., λ_3_ = −*δ*_2_. Finally, the vegetation-herbivore species found in the phase plane (*V* > 0, *S*_1_ > 0) gains an additional dimension, whose stability depends upon the critical value 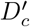,

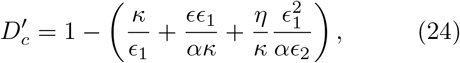

recall *κ* = *δ*/*μ*. For 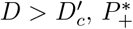 gains a stable direction in the full space. Comparing this to the stability condition (24) and the existence condition (11), it is easy to see that there is a parameter region where the fixed point is a repeller in the predator dimension and it turns to an attractor. This means that there is a region where tristability exists: the global extinction 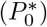, the global coexistence 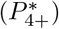 and the coexistence of vegetation and herbivours. This combined with the results from the previous section (the dynamics when *S*_2_ =0 should be preserved from the 2D model) indicate the herbivores-vegetation attractor can be either the fixed point 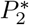 or the limit cycle (explained in the previous section) depending on the stability condition of 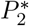 (see Eq. (11)).

The possible dynamics and the bifurcation boundaries of this system are displayed in Fig. 7. Panel A shows these dynamical regimes, which include coexistence and extinctions, in the parameter space (*κ, ϵ, D*). To illustrate qualitatively different dynamics we have selected different regions of this parameter space (see Fig. 7(a-f)): (a) bistable system with coexistence of all the species (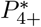 locally stable (LS)) and extinction (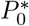 LS); (b) tristability with coexistence of all the species (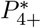 LS), herbivore-vegetation coexistence (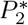 LS) and extinction (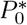 LS); (c) bistable state, herbivore-vegetation coexistence (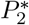 LS) and extinction (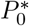 LS); (d) bistable state, herbivore-vegetation coexistence governed by a limit cycle (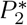 locally unstable) and extinction (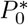 LS); (e) Bistable state (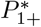 LS) and (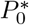 LS); and (f) global extinction (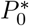 globally stable).Panels B and C show results for parameters *κ, μ, ϵ*, and *D* concerning existence of fixed points. Figure 8 displays an example of how different transient phenomena can concatenate. Specifically, the orbit shown first approaches the saddle-node ghost via oscillations and, after the extinction of predators, follows the spiral dynamics towards the full extinction. Note that the main delay is here caused by the ghost. The time series in Fig. 8 illustrate how this transient causes the chain of extinctions (similar to the extinction debt of metapopulations [50, 74, 75].

**FIG. 7:**
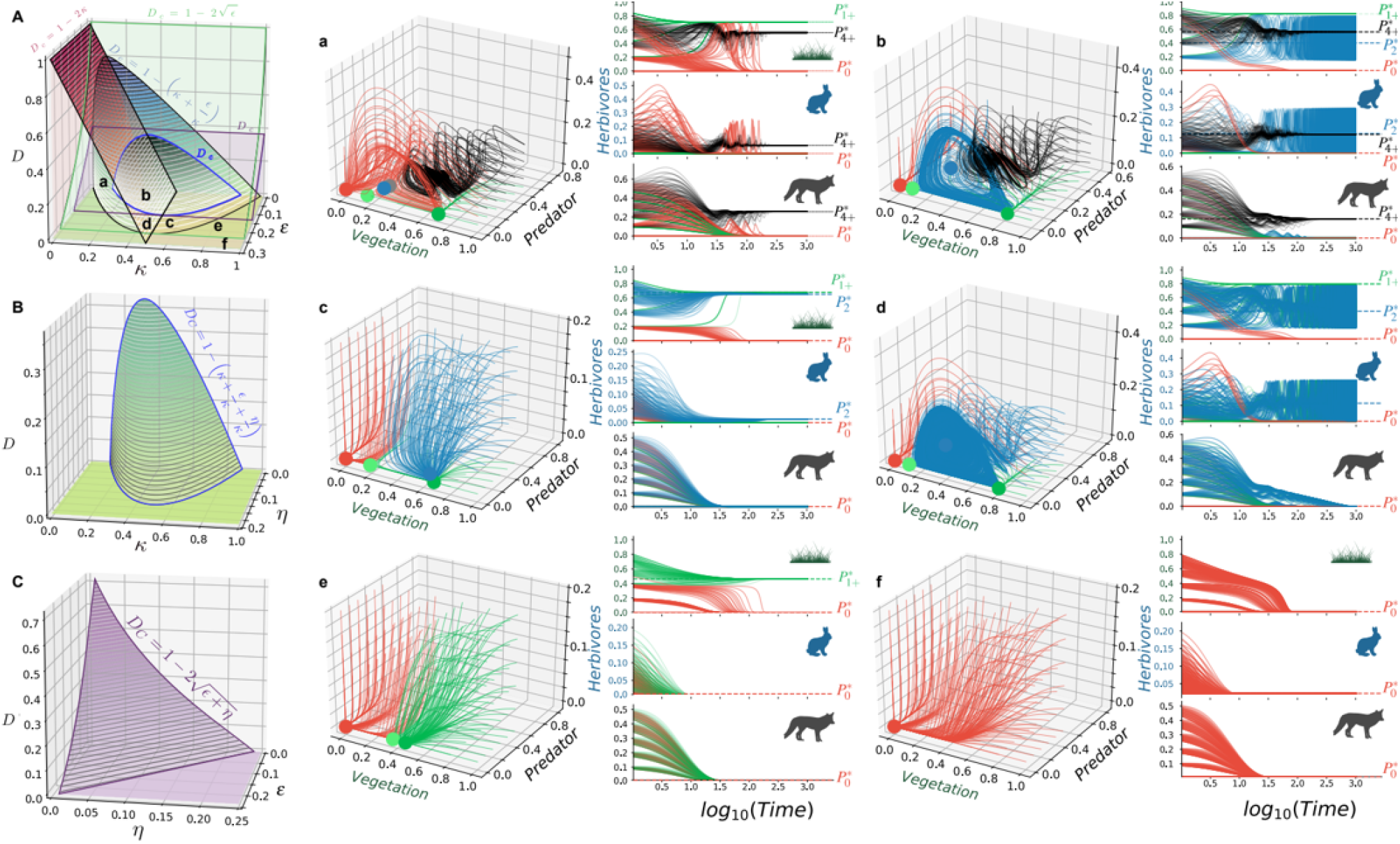
Dynamics, bifurcations, and transients for Eqs. (15)–(17). (A) Dynamical and stability boundaries in the parameter space (*κ, ϵ, D*), fixing *η* = 0.01. Relevant examples of dynamics are shown in the full phase space by means of phase portraits and time series for the regions indicated with small letters (a)-(f). (B) Minimal value of habitat destruction, *D*, needed to stabilise the dynamics at equilibrium 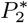, allowing for the coexistence between the vegetation and the herbivore species (boundary given by Eq. (24) here with *ϵ*=0.1). (C) Surface separating the full ecosystem coexistence governed by equilibrium 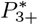, obtained from Eq. (23) (here setting *κ* = 0.35). The colours of the orbits in panels (a)-(f) indicate the state achieved by the system: full extinction (red), vegetation-only persistence (green); vegetation-herbivores coexistence (blue); full species coexistence (black). Notice that the system can be monostable (panel (f)); bistable (panels (c), (e), and (d); and tristable (panels (a) and (b)). The values of the equilibrium points are shown with the dots in the phase space and the dashed lines in the time series. In all panels: *α* = *μ* = *ρ* = *ϵ*_1_ = *ϵ*_2_ = 1.

**FIG. 8:**
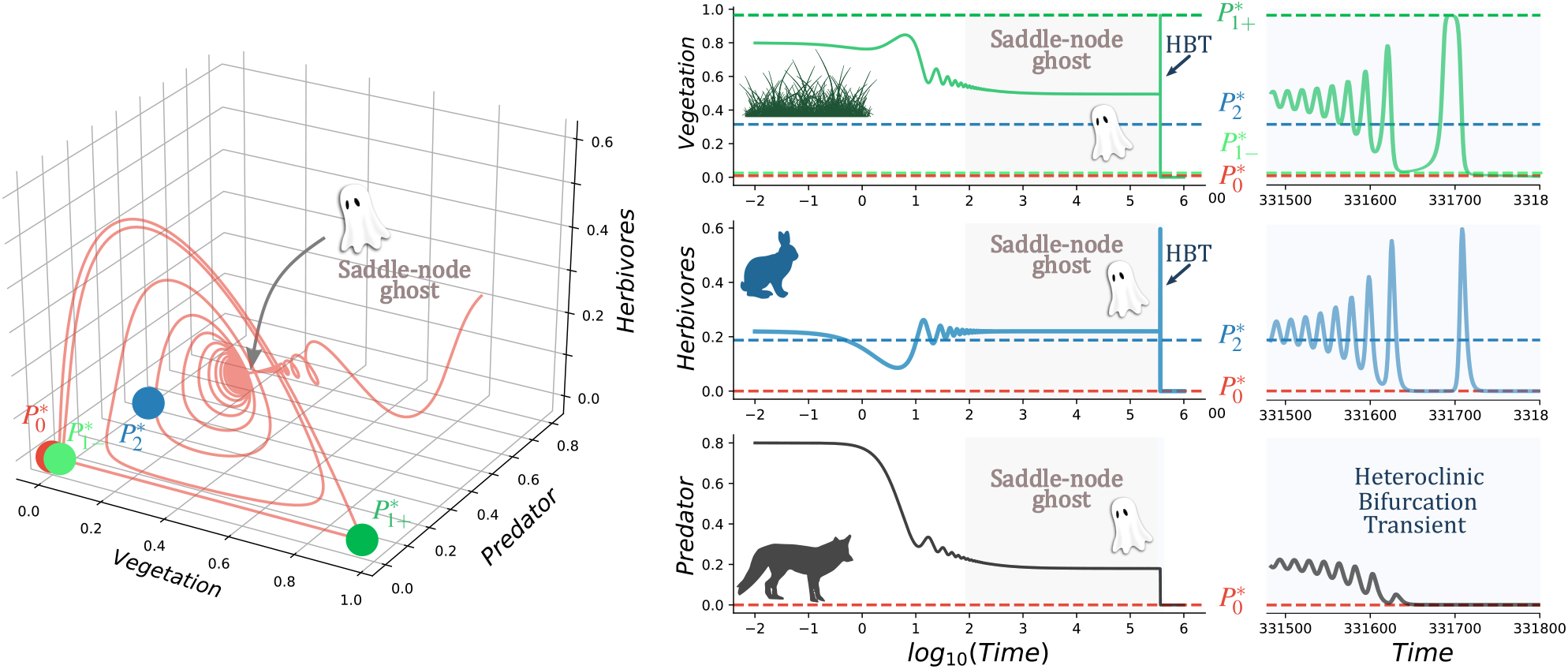
Example of a chain of transients towards extinction visualised in the full phase space and with time series. (Left) Typical path followed by an orbit starting far away from the region of the phase space where equilibria 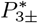 collided, leaving a ghost. Such a region sucks the orbit, which remains trapped for an extremely long time, and then spirals outwards experiencing the delays when they pass close to the saddle points 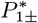 before going to the origin. (Right) Time series for vegetation, herbivores, and predators, with an initial oscillatory approach towards the ghost, a further delay settlet on the bottleneck region of the ghost (grey area), and the final large oscillations tied to the heteroclinic bifurcation transient (labelled as HBT). In all the analyses we use: *ϵ* = 0.025, *D* = 0.01, *α* = *μ* = *ρ* = 1.0, *δ* = 0.315, and *δ*_2_ = 0.220025.

### Larger trophic chains: future research

The models analysed in the previous sections can be extended to include further ecological complexity, as shown in Fig. 1(b). These simple models may also allow for a feasible investigation of the population dynamics on complex networks. Below, we propose two more models to be studied elsewhere. The first one adds an omnivore species that consumes the vegetation and predates on *S*_1_. The later includes a top predator consuming the omnivore species

### Model 4. Vegetation, herbivores and omnivores

This model considers the vegetation-herbivore species plus another species, *S*_3_, consuming the vegetation and predating the omnivore species. The model reads:

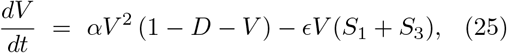

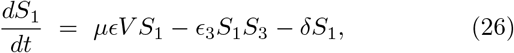

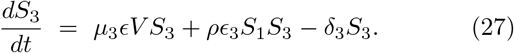

Here vegetation is consumed by the herbivore *S*_1_ and by *S*_3_, which grows proportionally to the term *μ*_3_*ϵ* > 0, here also with *μ*_3_ < 1. Omnivores are assumed to die proportionally to rate *δ*_3_ > 0.

### Model 5. Vegetation, herbivores, omnivores and a top predator

Here we consider the dynamics of the previous model adding a top predator, *S*_4_, that consumes the omnivore species, with dynamical equations:

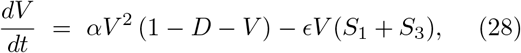

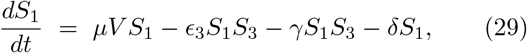

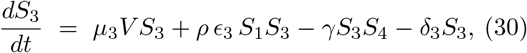

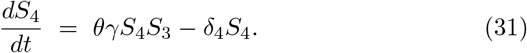

The reproduction rate of the top predator is proportional to *θ_γ_* > 0 and *θ* < 1, also assuming that they die proportionally to constant *δ*_4_ > 0.

## III. DISCUSSION

We have studied simple dynamical models for small trophic chains. Our main goal was to characterise how the addition of new species and thus new ecological interactions affected the dynamics, focusing on bifurcation phenomena and transients. We considered processes of facilitation at the level of primary producers, focusing on the impact of habitat loss, suggested as a key factor jeopardising species survival [47–50]. First, we have introduced a simple model for vegetation with facilitation recently studied [37]. This system is known to suffer a saddle-node bifurcation giving place to an abrupt tipping point as the habitat destruction overcomes a critical threshold.

The addition of a species consuming the vegetation introduces important changes in the dynamics, now allowing for the coexistence of both species and extinction of the omnivores or of the entire system. Here we identify a transcritical bifurcation separating the coexistence of both species and the extinction of the herbivores. The coexistence regime can be achieved via stable fixed point or via limit cycle. Interestingly, a global bifurcation driven by a heteroclinic bifurcation can induce a global extinction, in which the two species become extinct (a scenario dynamically analogous to the one found after the saddle-node bifurcation). The transient times and their dependence on bifurcation threshold have been determined. The transients after the heteroclinic bifurcation change in a discontinuous fashion as the habitat destruction approaches the critical value.

The previous system has been extended by adding a predator species consuming the herbivores. The most remarkable result of this model is that different phenomena responsible for transients can get coupled, thus enlarging transients. For instance, for some parameter regions, a ghost caused by a saddle-node bifurcation can get coupled with the oscillatory transient of the vegetation-herbivores system once the heteroclinic bifurcation has occurred. The coupling of different transients arising from different mechanisms may be relevant in theoretical ecology and may deserve further attention in future research.

Our results reflect the importance of so-called trophic downgrading [64]. The increase of hunting pressures on top predators (larger *δ*_2_ and *ρ* in our model), may cause a complete shift in dynamics, usually towards extinctions [76, 77]. During the transient, where all species remain, the reintroduction of predators could help in maintaining species and diversity, but effective changes may be introduced at the level of parameters (e.g. decreasing hunting of predators). The former strategy is the so-called rewilding [78–80]. Rewilding of wolfs in the Yellowstone National Park increased the complexity of the ecosystem, increasing the number of species living there and their abundances [81–83]. Once the wolfs became extinct, elks were out of control and they ate too much vegetation leading to lower *CO*_2_ capture and less food for other herbivores [83–85]. Moreover, the decrease in habitat space also led to simplifications of the trophic chain, even allowing only the persistence of producers. This is the so-called ecological meltdown, observed in some fenced areas in Africa [86–89]. For this case, what is being implemented are the formation of wildlife corridors [90–92], which increase the effective living area (meaning a decrease of habitat fragmentation parameter *D*). Notice that all the possible interventions should be done during the transients to ensure that the ecosystem is kept as similar as possible to the one found before multiple extinctions happened.

We note that the model explored here does not take into account complex functional responses [93–97], which introduce more realism to predator-prey dynamics. Despite this issue, our model includes the minimal interactions between the different tropic levels, allowing a clear identification of how different threads can push the system towards different kinds of bifurcations which have different properties (i.e. delayed extinction time) and signatures (changes in the temporal dynamics, see Fig. 6), similar to what is observed in real systems [31, 44, 98–100].

The presence of extremely long transients close to bifurcations (as shown in this article) opens the possibility that some ecosystems may be currently living in a ghost state, transitioning towards a less complex state [76, 77, 101] due to impact of human activity including deforestation, environmental contamination or defaunation. Anthropogenic impacts are making ecosystems to experience defaunation [102, 103], suffering extinction cascades due to trophic downgrading or ecological meltdown caused by ecosystem domestication (i.e. hunting top predators or cultivation fields expansion) [104–107]. These transients, where some species may be in a slow declining regime, or even in a false apparent stationary state, should provide opportunities for restoring ecosystems. Restoration strategies may be implemented during the transients [36, 108] to recover the maximum number of species and the complete ecosystem function (i.e. carbon and nitrogen fixation [109, 110]). More studies with real data should address this question. Also, future work may consider multiple species at each trophic layer [111], providing a closer approach to ecosystems as complex networks. Also, the impact of both intrinsic and extrinsic noise in the dynamics described in this contribution may be of interest.

## Acknowledgements

BV wants to thank Alan Hastings and Kim Cuddington for the invitation to contribute to the *Organized Oral Session on Transients in Ecology* at the Ecological Society of America in 2019. The authors thank Ricard Solé and Tomás Lázaro for useful discussions.

## Funding information

BV has been funded by the PR01018-EC-H2020-FET-Open MADONNA project and by the Botin Foundation, by Banco Santander through its Santander Universities Global Division. SV was supported by the Spanish Ministry of Economy and Competitiveness, grant FIS2016-77447-R MINECO/AEI/FEDER and the European Union. JS has been partially funded by the CERCA Programme of the Generalitat de Catalunya, by the MINECO grant MTM-2015-71509-C2-1-R and the Spain’s “Agencia Estatal de Investigación” RTI2018-098322-B-I00, and by a “Ramón y Cajal” contract (RYC-2017-22243). EF has been partially supported by the Spanish Government grant MTM2016-80117-P (MINECO/FEDER, UE) and by the Catalan Government grant 2017-SGR-1374.

## Conflict of interest

The authors declare that they have no conflict of interest.

## IV. APPENDIX

## A. Heteroclinic bifurcation

For the convenience of the reader, in this appendix we describe the key points of the heteroclinic bifurcation which takes place in Eqs. (6)–(7). It is similar to the Andronov-Leontovich homoclinic bifurcation, see for instance [112, 113]. Let us write our 2*D* system on a generic form as 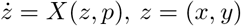, *z* = (*x, y*) or

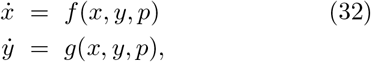

where *p* is a parameter. First, we present the setting of the heteroclinic bifurcation we deal with. The line {*y* = 0} is invariant and contains two equilibrium points *P*_1_ = (*x*_1_, 0), *P*_2_ = (*x*_2_, 0) of saddle type with 0 < *x*_1_ < *x*_2_. The position of *P*_1_, *P*_2_ may depend on the parameter *p*. Let λ_*i*_, *μ_i_* (here with *i* = 1,2), with λ_*i*_ < 0 < *μ_i_* be the eigenvalues of D*X*(*P_i_, p*). In general these eigenvalues depend on *p*. The unstable invariant manifold of *P*_1_, labeled *W^u^*(*P*_1_), and the stable invariant manifold of *P*_2_, labeled *W^s^*(*P*_2_), are contained in {*y* = 0}. Let *σ*_1_ be the corresponding heteroclinic connection between *P*_1_ and *P*_2_. Given *x_N_* ∈ (*x*_1_, *x*_2_), the branches of *W^s^*(*P*_1_) and *W^u^*(*P*_2_) in the region {*x* ≥ 0, *y* ≥ 0} cross the section {*x* = *x_N_*} at points *Q^s^*(*p*) and *Q^u^*(*p*) respectively. There exists a value of the parameter *p* = *p*_0_ such that *Q^s^*(*p*_0_) = *Q^u^*(*p*_0_). Let

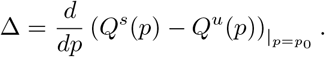

Let *σ*_2_ be the heteroclinic connection along the mentioned branches when *p* = *p*_0_. Now we assume two more specific conditions

a. 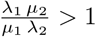,
b. Δ > 0.

Then, there exists a neighbourhood *U* of the heteroclinic cycle *σ_h_* = *σ*_1_ ∪ *σ*_2_ ∪ {*P*_1_, *P*_2_} and an interval (*p*_−_, *p*_+_) with *p*_0_ ∈ (*p*_−_, *p*_+_) such that for *p* ∈ (*p*_0_, *p*_+_) there exists a unique periodic orbit *γ_p_* of system (32) in *U* which is asymptotcially stable. Moreover, *γ_p_* tends to *σ_h_* when *p* → *p*_0_. For *p* ∈ (*p*_−_, *p*_0_) system (32) has no periodic orbits in *U*.

To prove the claim we introduce a return map close to *σ_h_* for values of *p* close to *p*_0_. Since our vector field is analytic there exists *C*^1^ local changes of coordinates *h*_1_ and *h*_2_ near *P*_1_ and *P*_2_, respectively, which (locally) linearize the system. Let *V*_1_, *V*_2_ be the domains of *h*_1_, *h*_2_. We take sections 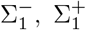 transversal to the flow in *V*_1_ which are preimages by *h*_1_ of the sections 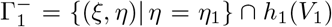 and 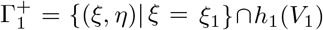 respectively, as shown in the image below.

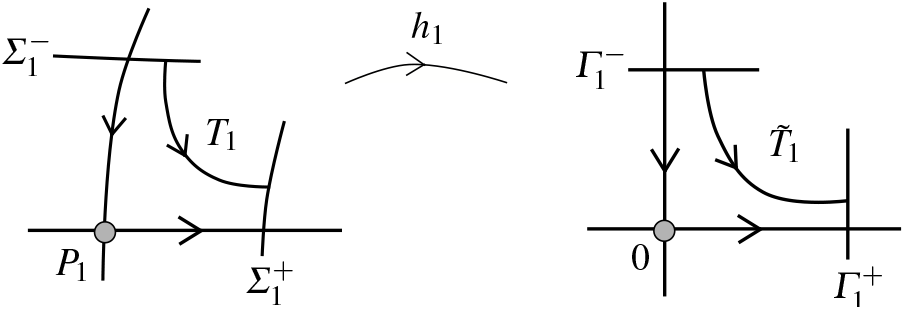

We define 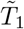 as the map which sends the coordinate *ξ* of the point 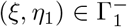 to the coordinate *η* of the point 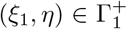 obtained as the intersection of the orbit of (*ξ, η*_1_) of the linear flow with 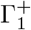.

The linearised system reads

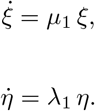

An easy computation from the general solution 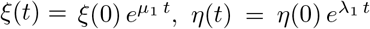, gives the time to arrive from (*ξ, η*_1_) to 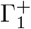:

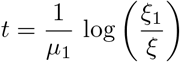

and then

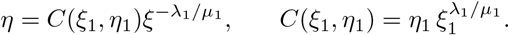

Going back by *h*^−1^ to (*x, y*) variables, using *ξ, η* as coordinates in 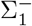 and 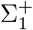 respectively, we have

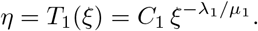

In a completely analogous way we define the map *T*_2_ from 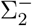 to 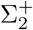, where 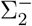 and 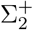 are the preimages by *h*_2_ of 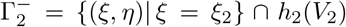 and 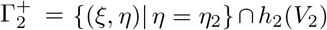 respectively. See the diagram below.

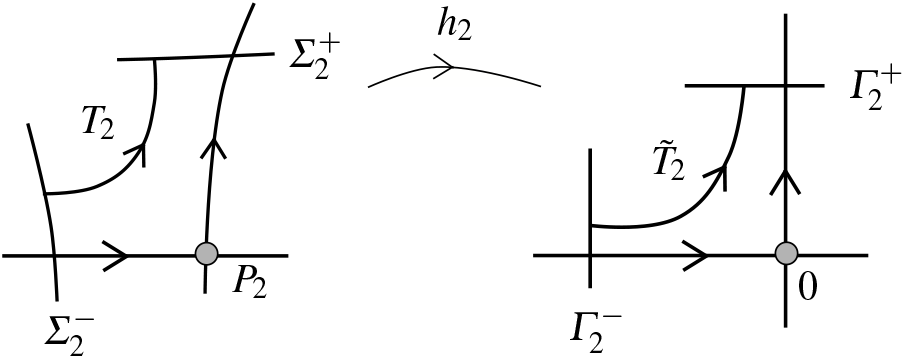

In local coordinates *u* in 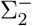 and *v* in 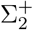

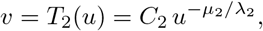

where *C*_2_ only depends on the chosen sections (and the parameter).

Also, we define 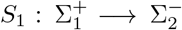 and 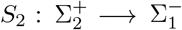 as regular Poincaré maps among the corresponding sections. They are analytic diffeomorphisms. Since the line {*y* = 0} is invariant for all values of the parameter, *S*_1_(0, *p*) = 0 and therefore

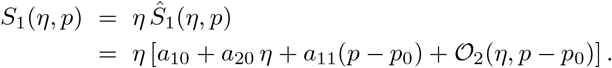

As for *S*_2_, we have *S*_2_(0, *p*_0_) = 0. The condition Δ > 0 implies

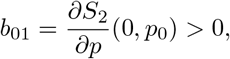

and then

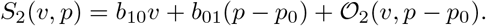

Since *S*_1_, *S*_2_ are diffeomorphisms we have *a*_10_, *b*_10_ ≠ 0. Moreover, since we are in the plane, the maps preserve orientation and therefore *a*_10_, *b*_10_ > 0. Then the return map 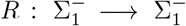 defined for *ξ* > 0 and *p* − *p*_0_ small as *R* = *S*_2_ ∘ *T*_2_ ∘ *S*_1_ ∘ *T*_1_ has the form

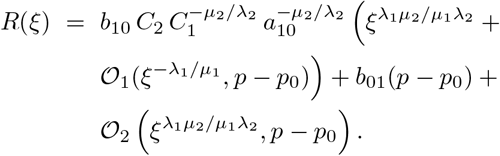

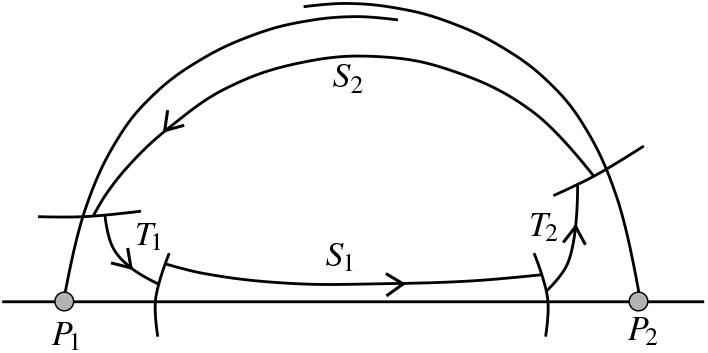

*R* is a perturbation of the 1*D* map

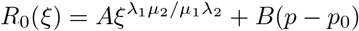

with *A, B* > 0 and λ_1_*μ*_2_/(*μ*_1_λ_2_) > 1. We display the graphs of *R*_0_ for *p* < *p*_0_, *p* = *p*_0_, and *p* > *p*_0_.

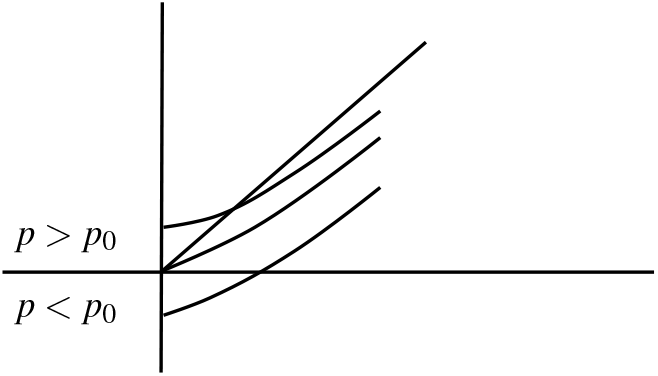

The claim follows directly from the interpretation of the image above. Indeed, notice that graph *R*_0_ cuts the diagonal close to *ξ* = 0 in and only if *p* > *p*_0_. The intersection point belongs to a periodic orbit. Moreover, since graph *R*_0_ crosses {*y* = *x*} from the upper to the lower part of the diagonal, the periodic orbit is asymptotically stable. Also, the fixed point tends to *ξ* = 0 when *p* tends to *p*_0_.

## B. Computation of the heteroclinic bifurcation value

Here we explain how the heteroclinic bifurcation value has been obtained by means of a numerical study of the global manifolds. The basic idea is to track when the STABLE MANIFOLD of the upper branch of equilibrium 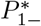 coincides with the UNSTABLE MANIFOLD of the upper branch of 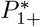. These two points have a HETEROCLINIC CONNECTION at bifurcation value. Before the bifurcation, a separation between the manifolds exists. After the bifurcation, a gap between the two manifolds will involve that all orbits go to the origin. Let us use the vegetation-herbivore model to illustrate the followed procedure (setting *μ* = 1 for simplicity and using *κ* = *δ*/*μ* = *δ*):

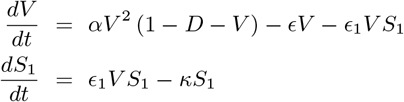

The basic procedure consists of:

1. Identify the equilibrium points that can get connected by a heteroclinic manifold. Here, there are three equilibria: the interior equilibrium (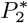, repeller focus) and two saddle points 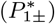.

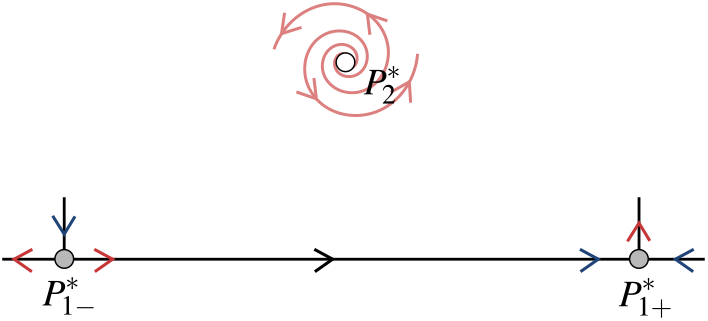
2. Calculate the eigenvectors 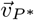 for each equilibrium point *P*\* that become heteroclinically connected i.e., the unstable manifold of one equilibrium becomes the same as the stable manifold of the other, here for 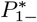 and 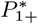. These eigenvectors read:

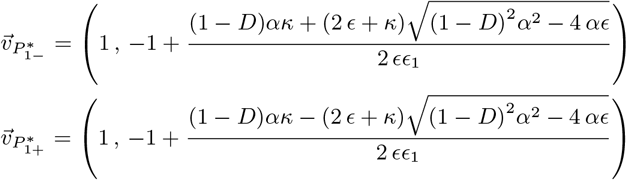
3. Compute the unstable manifold, *W^u^*. To do so, integrate the ODEs numerically using as initial condition 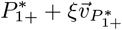 (using small *ξ*, here 10^−5^).

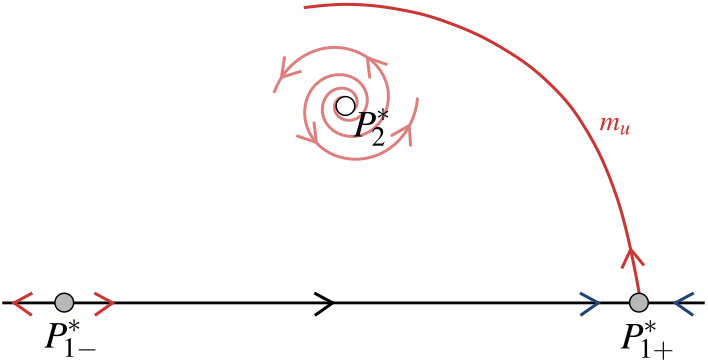
4. Compute the stable manifold, *W^s^*. To do so, we compute the solution of the ODEs numerically using as initial condition 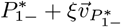. In this case, the solutions are computed in backward time. Since this manifold is stable, it attracts the orbits and the direction of the field needs to be changed to globalize the manifold.

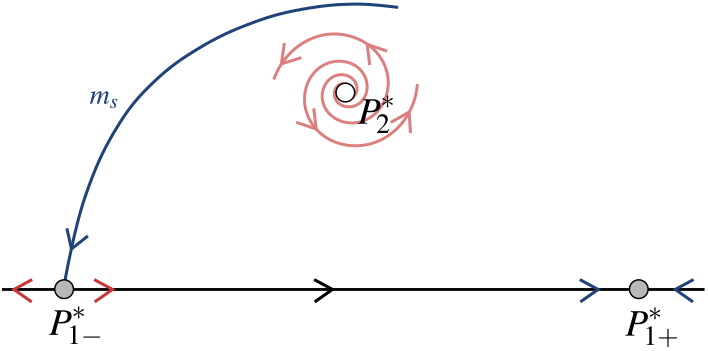
5. Chose a place (in the phase space) where the distance between the manifolds can be computed. Once the two manifolds collide they will collide simultaneously in all their trajectory (since they are heteroclinically connected). In the case shown in the main text, the value chosen was *V* = *κ*/*ε*_1_, the first time that this value is achieved. Let *m_u_* and *m_s_* be the intersection of *W^u^* and *W^s^* (for the first time) with the line *κ*/*ε*_1_ for first time.

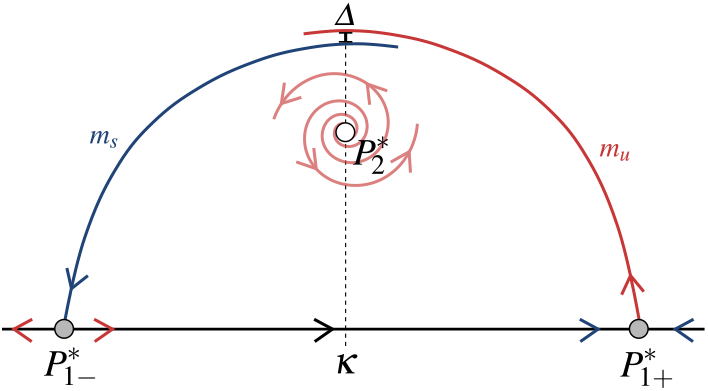
6. Compute the diference between the stable and unstable manifolds (Δ = *m_u_* − *m_s_*), while changing the parameter for which the critical value is desired. At some point, the distance will become negative, then the heteroclinic bifurcation had occured.

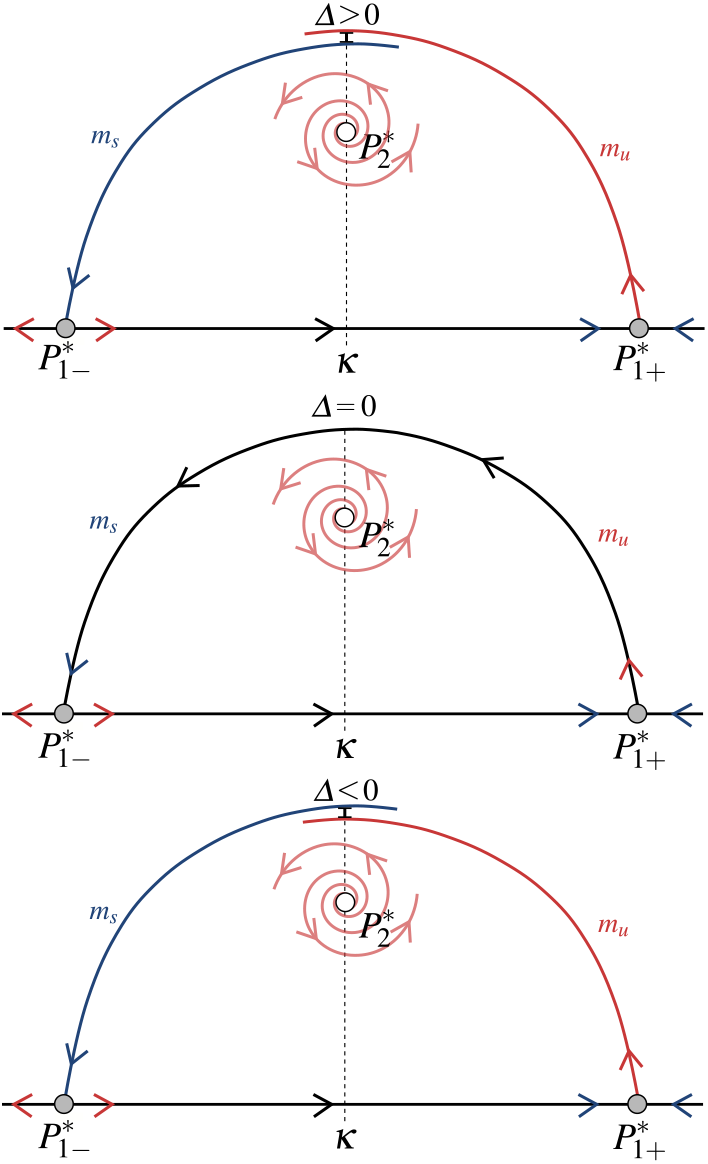 For the model studied in this article this procedure can be done in an easier manner. Knowing that the unstable manifold can only end-up going to the extinction fixed point (before the bifurcation), or to the interior fixed point (coexistence of vegetation and herbivores, after the bifurcation) the procedure can be simplified. The bifurcation value will be the parameter value for which there is a change between going to the extinction point or not.

## Notes

### Competing Interest Statement

The authors have declared no competing interest.

